# Floral signals evolve in a predictable way under artificial and pollinator selection in *Brassica rapa* using a G-matrix

**DOI:** 10.1101/675413

**Authors:** Pengjuan Zu, Florian P. Schiestl, Daniel Gervasi, Xin Li, Daniel Runcie, Frédéric Guillaume

## Abstract

**Background:** Angiosperms employ an astonishing variety of visual and olfactory floral signals that are generally thought to evolve under natural selection. Those morphological and chemical traits can form highly correlated sets of traits. It is not always clear which of these are used by pollinators as primary targets of selection and which would be indirectly selected by being linked to those primary targets. Quantitative genetics tools for predicting multiple traits response to selection have been developed since long and have advanced our understanding of evolution of genetically correlated traits in various biological systems. We use these tools to predict the evolutionary trajectories of floral traits and understand the selection pressures acting on them.

**Results:** We used data from an artificial and a pollinator (bumblebee, hoverfly) selection experiment with fast cycling *Brassica rapa* plants to predict evolutionary changes of 12 floral volatiles and 4 morphological floral traits in response to selection. Using the observed selection gradients and the genetic variance-covariance matrix (G-matrix) of the traits, we showed that the responses of most floral traits including volatiles were predicted in the right direction in artificial- and bumblebee-selection experiment, revealing direct and indirect targets of bumblebee selection. Genetic covariance had a mix of constraining and facilitating effects on evolutionary responses. We further revealed how G-matrices evolved in the selection processes.

**Conclusions:** Overall, our integrative study shows that floral signals, and especially volatiles, evolve under selection in a mostly predictable way, at least during short term evolution. Evolutionary constraints stemming from genetic covariance affected traits evolutionary trajectories and thus it is important to include genetic covariance for predicting the evolutionary changes of a comprehensive suite of traits. Other processes such as resource limitation and selfing also needs to be considered for a better understanding of floral trait evolution.

## Background

Understanding and predicting the evolutionary responses of phenotypes to selection remains a major challenge in evolutionary biology. This undertaking is not trivial because phenotypes are often complex traits co-evolving with other traits underlain by complex genetic architectures. Yet, understanding how such co-evolutionary units evolve under natural selection is important to understand how species may respond to changes in their environment. Flowers are complex organs with enormous diversity in morphology, color and scent, and thus represent a complex set of interrelated traits. These visual and olfactory components, which characterize the radiation of angiosperms, are recognized to evolve as a means of interaction with their biotic environment [1, 2]. One key driver, the pollinators, has been emphasized to be important for floral trait evolution since long [3, 4]. Yet, only a handful of studies have attempted to predict evolutionary responses of floral traits to pollinator selection [5–9]. Moreover, these studies only examined one or a few morphological traits at a time, whereas interactions of flowers with other organisms are typically mediated by a combination of traits of morphological and/or olfactory nature [10, 11]. A multivariate approach can, therefore, help to unravel the genetic architecture of floral traits and predict their joint evolution.

A great number of empirical studies have documented significant heritability and genetic (co)variance of diverse floral traits [12–15], as well as phenotypic selection acting on them [16–26]. Among those traits, floral scents have started drawing more and more attention. Floral scents are usually highly variable and diverse on all taxonomic levels [27], and many studies have documented natural selection on scent [22–25, 28, 29]. We have shown that scent phenotypic variation has a significant heritable genetic component in fast-cycling *Brassica rapa* (20% – 45%, [15]). In the same species, Gervasi and Schiestl [25] showed in a laboratory experiment that bumblebee pollinator selection resulted in taller plants with larger flowers and increased amounts of several floral scents, presumably used as visual and olfactory signals for more nectariferous plants. However, not all trait changes, or the absence of change, could be explained from the estimate of the selection intensities acting on them [25]. Phenotypic trait responses to selection are known to depend on the pattern of genetic variance-covariance among them [30, 31]. In particular, traits that are genetically correlated because of a shared genetic basis (i.e., pleiotropic genes) will indirectly respond to selection on linked traits, which may mask the effect of direct selection on them. Therefore, the direct targets of pollinator selection cannot be well characterized unless the selection responses are decomposed into their direct and indirect components. This is best done using a multivariate quantitative genetics framework [30, 32].

Quantitative genetics theory provides a means to make such evolutionary predictions in the form of the multivariate breeder’s equation (or Lande’s equation), Δ**z** = **G*β** [30]. Lande’s equation predicts the per-generation change in a set of quantitative traits in a population (Δ**z**) as the product of their genetic variance-covariance matrix (G-matrix) with the vector of selection gradients acting on them (**β**). One key insight here is that traits may deviate from their predicted changes under direct selection (**β**) because of indirect selection pressures caused by selection on the other traits and their genetic correlation with them. In other words, traits may deviate from their expected evolutionary trajectory given by **β**, and thus be constrained by genetic correlations with traits under a different set of selection pressures [33, 34]. The components of Lande’s equation can be estimated from phenotypic and individual pedigree relationship data in an experiment by using the classical tools of quantitative genetics [31, 35]. Moreover, this multivariate approach can help distinguish between the direct and indirect targets of pollinator selection. This distinction can be made when comparing the direct and indirect components of a trait’s predicted evolutionary response with its observed selection response. The direct component is obtained when multiplying the diagonal elements of a G-matrix (*G_ii_*, the additive genetic variance of the traits) with the **β** vector, which holds, for a single trait *i*: 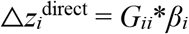, while its indirect selection response is the product of the off-diagonal elements of **G** (genetic covariance: *G_ij_*) with **β**, summed over all traits *j* ≠ *i*: 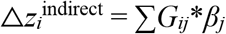. The total response is the sum of these two components. Traits that have the same direction of observed and direct selection responses can be considered as direct targets of selection. In contrast, indirect targets would be traits with same direction of their observed and indirect responses but opposed direct component of the response. In sum, if the total predicted response of a trait is opposed to its direct component, or much smaller, evolution of that trait will be said evolutionarily constrained by the genetic correlation among the traits.

In this study, we predict floral traits evolution under artificial and pollinator selection. We tested how the evolutionary trajectory of a trait would be affected (constrained/enhanced) by genetic correlations among traits by dissecting the total responses to selection into direct responses (caused by direct selection on target traits) and indirect responses (caused by correlated responses through genetic covariance). Moreover, we also assessed the evolution of genetic architectures (G-matrices) throughout the artificial selection processes. We used data from two forward-in-time experimental evolution experiments that documented genetic covariation and evolutionary responses in floral traits of fast cycling *Brassica rapa* plants. The G-matrix of the plant population was estimated from a three-generation bi-directional artificial selection experiment on plant height [14]. In that study, tall- and short-plants were selected artificially for building the two directional lines, plus randomly selected plants for an additional control line. Four morphological floral traits and 12 floral volatiles were measured in each generation. Control lines in this experiment were used to estimate the G-matrix. The selection gradient β was estimated in four evolutionary experiments: two for the tall- and short-selection lines in the artificial selection experiment mentioned above [14]; the other two from a 9-generation pollinator selection experiment [25]. The pollinator selection experiment was carried out with bumblebees and hoverflies as the selection agents separately. The same set of floral traits were measured, and the parental plants were from the same seed bank as in the artificial-selection experiment.

## Results

### Predictions in the artificial selection experiment

The response of plant height, the direct and only target of artificial selection, was correctly predicted in both treatments (Fig. 1). Given that all other traits are positively correlated with plant height (see Table S4), their indirect responses are predicted positive for tall lines and negative for short lines. However, observed responses of the flower size traits (petal width, PW; petal length, PL; and flower diameter, FD) were correctly predicted only in short lines, while the predictions to increase in tall lines mismatched observations (Fig. 1). The correlated responses of floral volatile organic compounds (VOCs) were predicted well in half of the traits in both treatments (Fig. 1). The predictions for the other half of the VOCs were not significant because HPD intervals of predicted values overlapped with zero (Fig. 1).

**Figure 1.**
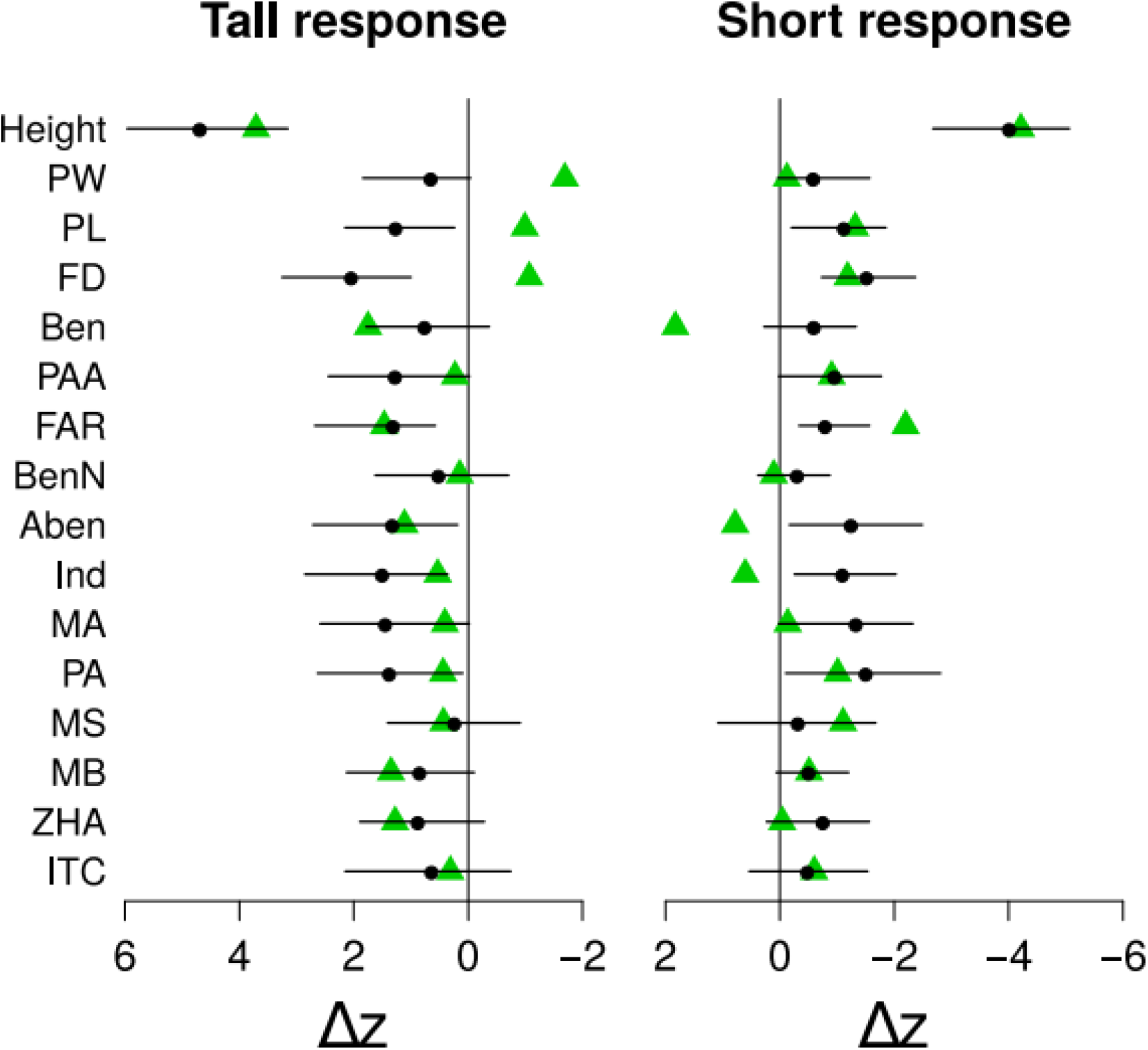
Predicted and observed responses of various traits to artificial selection. Green triangles are the observed changes. Black dots are the predicted selection responses. The solid horizontal lines indicate the 95% HPD interval of the predictions. Both predicted and observed changes were scaled by the phenotypic standard deviation of the trait. Sample sizes: plant height: 600; flower size traits (PW, PL, FD): 581; volatiles: 579. Trait abbreviations: Plant height (Height), Petal width (PW), Petal length (PL), Flower diameter (FD), Benzaldehyde (Ben), Phenylacetaldehyde (PAA), α-Farnesene (FAR), Benzyl nitrile (BenN), 2-Amino benzaldehyde (Aben), Indole (Ind), Methyl anthranilate (MA), phenylethyl alcohol (PA), Methyl salicylate (MS), Methyl benzoate (MB), Z-(3)-Hexenyl acetate (ZHA), 1-Butene-4-isothiocyante (ITC).

### Predictions in the pollinator-selection experiment

In the bumblebee treatment, all but one VOC (methyl salicylate, MS) increased during the experiment (Fig. 2A; Table S6). Our predictions somewhat overestimated the evolutionary changes of morphological traits, especially height and FD. Responses of scent compounds were correctly predicted in 7 VOCs, although only 4 of them (alpha-farnesene, FAR; indole, Ind; methyl anthranilate, MA; and methyl benzoate, MB) were significantly different from zero (Fig. 2A; Table S6). Observed responses were significantly larger than their predictions in phenylacetaldehyde, PAA; benzyl nitrile, BenN; 2-amino benzaldehyde, Aben, and phenylethyl alcohol, PA (Fig. 2A; Table S6).

**Figure 2.**
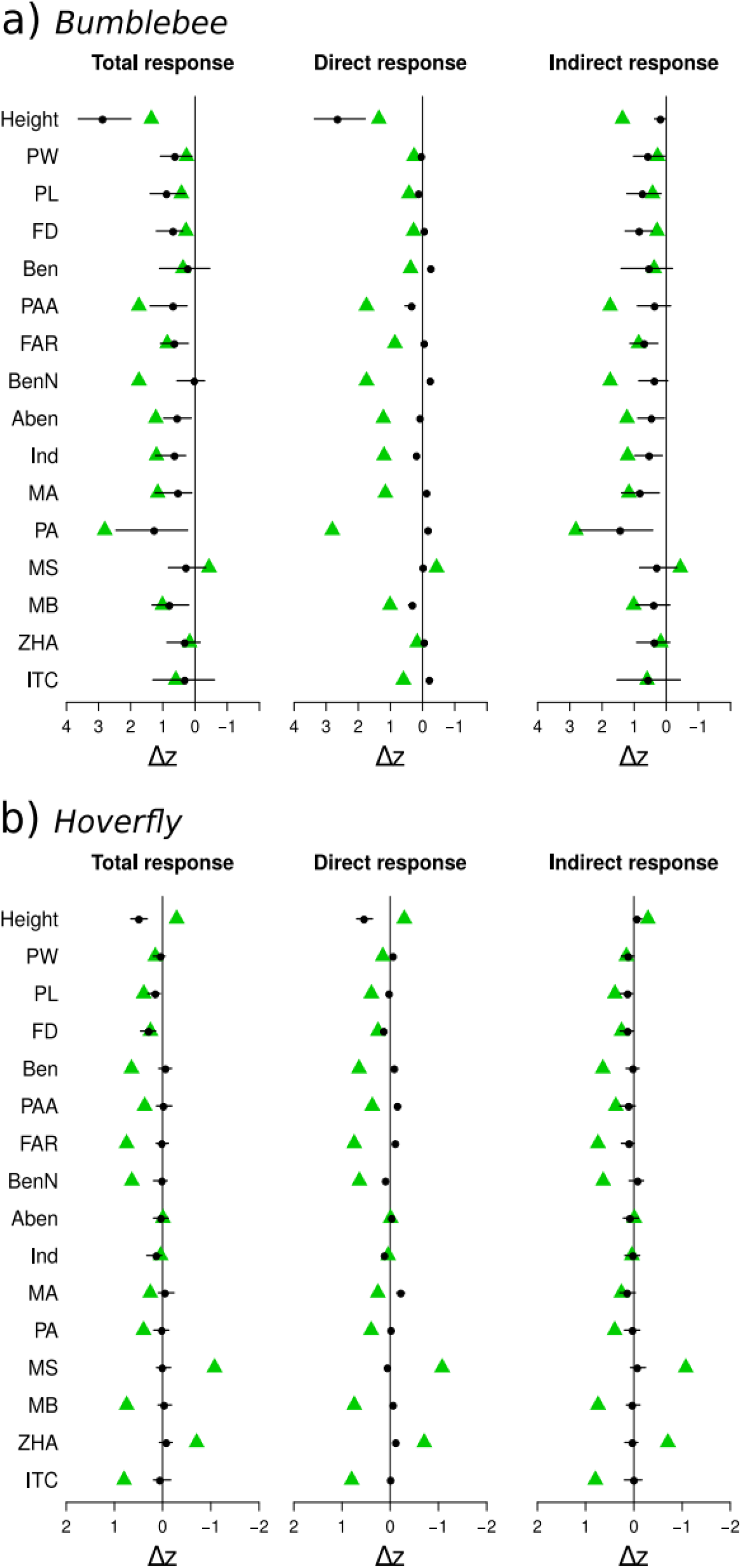
Predicted and observed responses of various traits to pollinator selection, in the bumblebee (a), and hoverfly (b) experiments. The total predicted response of each trait is decomposed into its direct and indirect components (see text). Green triangles are the observed changes. Black dots are the predicted selection responses. The solid horizontal lines indicate the 95% HPD interval of the predictions. Both predicted and observed changes were scaled by the phenotypic standard deviation of the trait. Sample sizes pollinator selection: plant height: 524, flower traits: 525, volatiles: 414. Same trait abbreviations as in Figure 1.

The response decomposition analysis showed that most direct responses were much smaller and in the wrong direction relative to the observed responses. The direction was correctly predicted as positive for plant Height, PW, and PL among morphological traits, and PAA, Aben, Ind, and MB among VOCs (Fig. 2A). Only plant height’s direct response overestimated its observation, while all others underestimated theirs. In contrast, the indirect components of the predicted responses were in the right direction for all traits (but MS), and of the right magnitude for five VOCs (Ben, FAR, MA, ZHA, and ITC) plus PW and PL. Therefore, the VOC and petal size responses are mostly indirect responses to selection on other traits.

In the hoverfly treatment, trait responses were small compared to the bumblebee treatment (Fig. 2B). Evolutionary predictions were mostly not different from zero except for FD and plant height, although the height response was opposed to its prediction. Of the 12 VOCs, only Ind and Aben responses were within their prediction’s HPD intervals, although Aben’s response is not different from zero (Fig. 2B; Table S6). From observed and predicted responses of morphological traits, hoverfly selection seemed to have targeted flower size more than plant height. Decomposition of the trait responses showed the same overall pattern as in the bumblebee experiment but with much smaller predicted than observed responses even for the indirect components. Overall, observed and predicted changes were larger in the bumblebee than the hoverfly treatment (Fig. 2B; Table S6) in agreement with the weaker selection gradient for hoverfly than bumblebee selection (Table S2).

### Effects of genetic covariance on predicting evolutionary trajectories

We measured the overall constraining effect of genetic co-variation on the response to selection by comparing the angle **θ** between the selection response vector (Δ**z**) and the first PC of **G** (PC1, or ***g***_max_, see methods) with the angle **γ** between Δ**z** and the selection gradient (**β**). In the tall and short artificial selection experiments, the trait responses were strongly aligned with ***g***_max_, with **θ** angle of 12.5 degree (95% HPD: 9.2, 16.6) and 11.2 degree (95% HPD: 7.2, 14.7), respectively. Given the close association of ***g***_max_ with the first trait axis (height) (Fig. 3c) and thus with the selection gradients under artificial selection, the angle **γ** between Δ**z** and **β** is 9.9 and 8.9 degree in tall and short, respectively, which are within the 95% HPD of **θ** in both cases. In contrast, under pollinator selection, Δ**z** is more aligned with ***g***_max_ than **β**, with **θ** of 26.9 degree (95% HPD: 22.3, 33) and 60.8 degree (95% HPD: 57.1, 63.5), when compared to **γ**, equal to 66.5 and 89.2 degree for bumblebee and hoverfly treatments, respectively.

**Figure 3.**
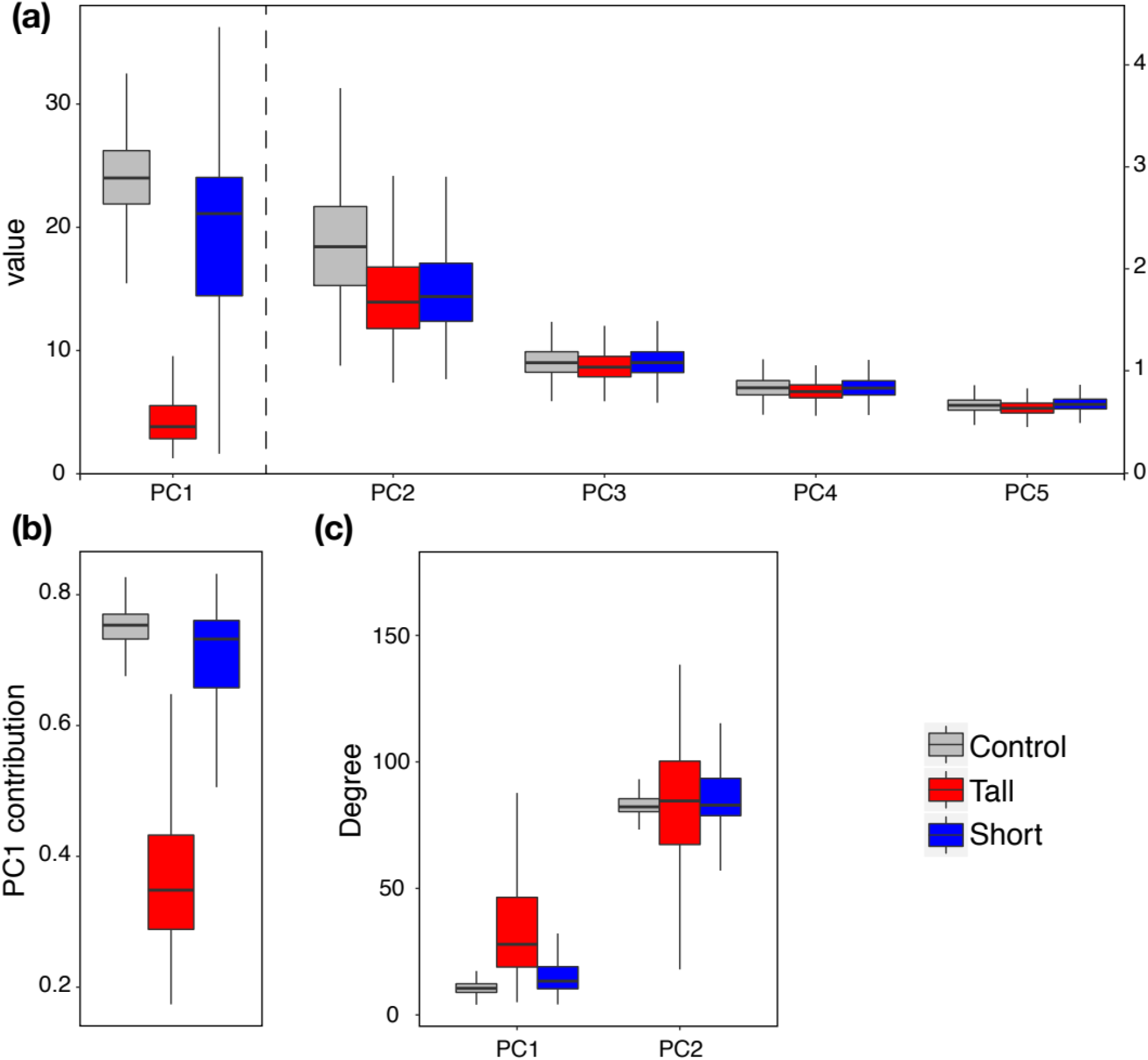
Comparison of the size and orientation of the major and five first eigenvectors (PCs) of the **G**-matrices in the artificial selection experiment. **(a)** Distribution of the eigenvalues (size) of each PC of the three **G**-matrices in the control (grey), tall (red), and short (blue) artificial selection experiments. The scale of the *y*-axis is on the left for PC1 and on the right for PC2-5. **(b)** Contribution of PC1 to the total variation in the 16 traits, measured as the size of PC1 relative to the sum of all PCs. **(c)** Angle of the first and second PC with the first trait axis (height) in degree. In all cases, variation of all variables stems from the posterior distribution of each **G**-matrix estimated with MCMCglmm (see Methods and Supporting information).

### Evolution of the G-matrix during artificial selection

By examining the G-matrices of the three lines in the artificial selection experiment (**G**_control_, **G**_tall_, and **G**_short_), we found a drastic decrease of the additive genetic variance of height in the tall line, with an estimate around 2.8 cm^2^, compared to the short line, which remained as high as in the control line around 23 cm^2^. This resulted in a large decrease of the contribution of ***g***_max_ (PC1) of **G**_tall_ to the total variance relative to **G**_control_ and **G**_short_ (see Fig. 3a-b, Table S5). The orientation of ***g***_max_ also changed in **G**_tall_, with reduced alignment with the height axis (Fig. 3c). The other eigenvalues and eigenvectors are, however, more constant across lines (Fig. 3a). For instance, the second eigenvector (PC2) is more consistently orthogonal to the height trait axis in the three **G**-matrices (Fig. 3c).

To further compare G-matrices, we used two different approaches from the toolkit of G-matrix comparisons, the random skewers and CPC approaches (Roff et al. 2012, see Methods). Using the random skewers method, we found strong correlations of the mean selection response among matrices, larger than 70% for all three comparisons, although not significantly so between **G**_tall_ and **G**_short_, and very strong similarity between **G**_control_ and **G**_short_ (Table 1). The three G-matrices thus shared a significant portion of their structure. **G**_control_ would predict selection responses similar to **G**_short_ and to a lesser extent to **G**_tall_. Further analysis of the similarity of the size and orientation of the eigenvectors of the G-matrices in the hierarchical analysis (CPC) confirmed the similarity in shape between **G**_control_ and **G**_short_ and the dissimilarity of **G**_tall_ with **G**_control_, and with **G**_short_ to a smaller degree (see Table 1). The G-matrix in the tall lines thus evolved more than in the short lines mostly because of the change in the genetic variance of plant height. **G**_short_ remained closer to the starting G-matrix (**G**_control_) over the course of the experiment.

**Table 1.** Comparisons of the three G-matrices using random skewers and hierarchical analyses. (see also Table S3, S4). The random skewers section reports the mean correlation among response vectors of two **G**-matrices subject to the same set of 10,000 random selection vectors. The hierarchical analysis reports the *P*-values to reject the hypotheses of equality, proportionality, or common principal components (CPC) in favor of *unrelated* matrices. The *P*-values are obtained by randomization (see Methods).

| Paired G | Random Skewers |  | Hierarchical |  |  |
| --- | --- | --- | --- | --- | --- |
|  | Mean correlation | *P*-value | Equal | Prop. | CPC |
| **G**<sub>control</sub> - **G**<sub>tall</sub> | 0.734 | < 0.01<sup>a</sup> | < 0.005 | < 0.005 | 0.002 |
| **G**<sub>control</sub> - **G**<sub>short</sub> | 0.987 | < 0.002<sup>b</sup> | 0.23 | 0.23 | 0.32 |
| **G**<sub>tall</sub> - **G**<sub>short</sub> | 0.722 | < 0.21<sup>a</sup> | 0.19 | 0.19 | < 0.05 |

<sup>a</sup>: left tail; <sup>b</sup>: right tail

### Evaluation of the estimation of the **G**-matrices

Permutation tests of G-matrices were conducted to examine whether our G-matrix estimates captured the meaningful biological structure of the data. The results revealed that the majority of the genetic covariance elements (101 out of 120) and additive genetic variances (14 out of 16) in **G**_control_ were significantly different from zero at the level of FDR < 0.05 with 500 permutations and after correcting for multiple testing (false discovery rate: Benjamini & Hochberg, 1995). In **G**_tall_, 11 variance and 55 covariance elements were significant, and 15 and 72 elements, respectively, in **G**_short_ (Table S4), at the same FDR level. Furthermore, variance estimates had much narrower 95% HPD intervals than covariance estimates (from their posterior distributions, results not shown), as evident in the size of HPD intervals of the direct and indirect components of the selection responses (see Fig. 2).

## Discussion

Total evolutionary trait responses are made of direct and indirect responses. Evolutionary constraints emerge when the two oppose each other. However, constraints may evolve when the genetic variance and covariance among traits, that is the elements of the G-matrix, change over time, caused by selection or other evolutionary forces. It is thus important to evaluate the structure of the G-matrix and its evolution when trying to understand the effects of selection on multiple phenotypic traits. Moreover, being able to compare predicted and realized trait responses allows for a better understanding of the relationship between selection and genetic constraints. In this study, we could predict the evolutionary response of floral traits subject to two types of selection pressures by combining estimates of the ancestral G-matrix of the traits with estimates of the selection gradients acting on them. Importantly, we found that predictions based only on the direct trait responses to selection failed to predict the observed responses and that the observed responses were biased towards the line of least genetic resistance (***g***_max_) of the G-matrix. The pattern of genetic covariation among traits thus strongly affected the outcome of selection in the artificial and pollinator selection experiments. Although this pattern of trait covariation can change during evolution, we further showed that using an ancestral G-matrix, here estimated in the control lines, can lead to accurate evolutionary predictions over just a few generations. This approach allowed us to better understand how pollinators, the selective agents, interact with the complex set of floral traits composed of floral scent and morphology and may influence their evolution.

Overall, bumblebee selection was in favor of taller plants with bigger petals and increased concentration of certain odor volatiles, most notably indole (Ind), phenylacetaldehyde (PAA), 2-amino benzaldehyde (Aben), and methyl benzoate (MB) [see also 25]. Those traits had a significant positive selection gradient in the same direction as their observed response making them candidates for direct targets of bumblebee selection. The indirect components of the responses were also all positive, enhancing the total predicted responses, sometimes leading to overshooting of the observed responses. Because of the largely positive genetic correlation of most floral traits with height, it is not surprising to observe positive total selection responses of most traits, and our evolutionary predictions match well with that pattern. However, our analysis revealed that some of those responses may be maladaptive because opposed to the selection gradient acting on them (e.g., Ben, BenN, PA, and ITC, see Figure 2, Table S6). This suggests that bumblebees tended to dislike flowers with increased concentration of those volatiles, whose increased observed amounts were caused by indirect selection on height and other positively correlated traits under positive selection. Their positive, non-adaptive responses thus point to the existence of strong evolutionary constraints acting on them. Overall, knowledge of the selection gradient, the G-matrix and selection response of the traits showed that they evolved in a direction biased towards ***g***_max_, the “line of least resistance” [33], which constrained the evolutionary response away from the selection gradient, although the selection responses of some traits were enhanced by trait covariation.

In contrast to predictions in the bumblebee-pollinated plants, the ones in hoverfly-pollinated plants were largely not different from zero or incorrect. Only changes in flower size were correctly predicted. The observed changes were also not consistently in the same direction, implying that an evolutionary response along one major axis of overall positive trait co-variation is not likely, at least when estimating the co-linearity of the response vector with ***g***_max_ of **G**_control_. Instead, the observed changes are more consistent with very weak selection and altered patterns of trait covariation. Indeed, in the hoverfly-selection experiment, a separate study found very little adaptive evolution in plant traits with the exception of strongly increased autonomous selfing [25]. Thus, increased selfing and the associated reduction of genetic variation [37], possibly altered the G-matrix, leading to the low accuracy of our predictions and the reduced efficiency of pollinator-induced selection. Previous studies in bottlenecked insect populations have shown that rapid changes in the G-matrix are expected in inbred populations [e.g., 38, 39].

We observed further discrepancies between our evolutionary predictions and observed responses that need to be examined. In particular, the responses of the morphological traits in the artificial selection for tall plants did not show the expected increase of flower size but instead showed a decrease, despite the positive genetic correlations of flower size with plant height (see **G**_control_ in Table S4), which remained positive during selection (see **G**_tall_ in Table S4). It thus cannot be caused by an evolutionary change of the sign of the genetic correlations with height. Instead, this selection experiment may have revealed an underlying resource allocation trade-off masked by the apparent positive genetic covariation between plant height and the size of the reproductive organs. This is reminiscent of classical theory on the effect of variation in resource acquisition and allocation on fitness components [40–42], which states that a positive correlation between fitness components can be observed despite an underlying trade-off when individuals vary more in the acquisition than in the allocation of their resources. Variation in resource acquisition among the genotypes may have been pre-existing in the base population of *B. rapa*, and lead to the observed positive correlation between traits pertaining to two fitness components, plant reproduction for flower size traits, and plant somatic growth for plant height. However, the observation of a negative correlated response of flower size to selection for reduced plant height in the low artificial selection experiment is more in line with a positive correlation between height and floral morphology. This may be caused by unconstrained allometries when selecting for smaller plants where resource limitation may be less stringent than when selecting for taller plants. Finally, the large overshooting of the predictions of the response of plant height in the bumblebee and hoverfly experiments is probably due to an overestimation of its genetic variance, although we cannot test for this hypothesis because we are missing an estimate of the G-matrix in that selection experiment.

### The role of genetic covariance in adaptive evolution

Our results are in line with the established expectation that genetic covariance can influence traits’ evolutionary responses by constraining or augmenting their response to selection depending on the relative signs of genetic covariances and selection gradients [30, 32, 43]. This expectation has been rarely directly tested with experimental evolution as we did here [see also 44]. More commonly, empirical studies use estimates of contemporary selection gradients and G-matrices to evaluate the potential for evolutionary constraints, which are present in some cases (e.g., [45–48]) but not in others (e.g., [34, 49, 50]).

The relevance of predictions of evolutionary constraints depends on the constancy of patterns of genetic variance-covariance over time. Our study shows that constancy cannot be assured when selection strongly reduces the genetic variance of a trait, as during artificial selection for taller plants (see also [44, 51, 52]). Yet, using **G**_control_ as an estimate of the ancestral G-matrix allowed us to make correct evolutionary predictions of the direction of selection responses in most cases. Had we used **G**_tall_ in the tall selection experiment, we would have badly underestimated the selection response of plant height and floral scents (results not shown). This illustrates two important points concerning the evolutionary significance of the structure of the G-matrix. First, changes in **G** can happen quickly, over just a few generations, and we have illustrated a rapid change in trait variance caused by selection. Second, despite those changes, estimation of **G** is still useful to make predictions of future trait changes over few generations. This can be useful to predict evolution and adaptation under rapid environmental changes, for instance, because the state of the G-matrix before a change in selection pressures will strongly influence the resulting evolutionary trajectory of a population, as we have shown here.

The evolutionary significance of the structure of the G-matrix is still debated, especially regarding the interpretation of the constraining effects of the main eigenvectors of **G** (especially ***g***_max_). The debate, however, mostly crystallized on inferences of past evolutionary constraints from contemporary estimates of trait variance-covariance patterns. The retrospective use of **G** is questionable knowing how evolutionarily labile are patterns of variance-covariance, an important caveat already emphasized by Turelli [53]. Indeed, many processes may affect the evolution of trait variance and covariances because they depend on variation in allele frequencies in a population. As such, genetic drift [54] and fluctuating selection [55], have been shown to reduce the stability of the G-matrix, while migration [56], correlational selection [54], and mutation [54, 57] can improve its stability [reviewed in 58]. Those changes thus make retrospective use of **G** at the least dangerous, unless its long-term stability can be determined. Prospective use of **G** is potentially less sensitive to such variations when predicting short term selection responses. Our analysis provides a good illustration of the prospective versus retrospective usage of a G-matrix when considering the changes in **G**’s structure between **G**_control_ and **G**_tall_ and the respective predictions and inferences we can make from them.

## Conclusion

Our study showed that even highly plastic chemical traits such as floral scent, can be successfully included into predictive models of floral trait evolution. Even more so, we show that a complementary set of traits is important to consider, because pollinator selection acts on multiple traits, and genetic correlations link them in their evolutionary response. In the future, improved sampling and analysis techniques may allow the standard inclusion of a large set of traits and large sample sizes into evolutionary studies. Larger sample sizes may allow for more accurate predictions by incorporating the dynamics of G-matrix evolution over multiple generations. In addition, more assessments of selection on those traits in nature by specific groups of interacting organisms [21, 23, 24, 29, 59] may further improve our ability to predict evolutionary changes in the face of environmental change in natural habitats.

## Materials and methods

The workflow of the main analysis procedures is summarized in Fig. S1.

### Plant species and focal traits

In our experiment, we used the rapid cycling *Brassica rapa* L. (syn. *B. campestris*: Brassicaceae) from the Wisconsin Fast Plants™ Program (Carolina Biological Supply Company, Burlington, NC, USA), selected for short generation time, rapid seed maturation, absence of seed dormancy, small plant size and high female fertility [60]. *Brassica rapa* is a self-incompatible species with a generalized pollination system (e.g. bees, syrphid flies and butterflies as pollinators). The line used needs only ca. 35 to 40 days to complete a life cycle and maintains sufficient genetic variability for selection experiments [14, 15, 61, 62].

Our analysis includes a total of 16 traits with 12 floral volatile organic compounds (VOCs), and 4 morphological traits (plant height, petal width, petal length, and flower diameter). The measurement methods were described in detail in Zu & Schiestl [14]. Floral VOCs were collected from at least four freshly opened flowers per plant at a flow rate of 100 mL per min for 3 hours. Floral VOC amounts were standardized in amounts per flower per liter sampled air, and log transformed (by ln(x +1), x being the raw value) prior to statistical analysis to approach normal distributions. Scent collection and analysis details can be found in Supporting information. The whole experiment was conducted at the Botanical Garden of the University of Zürich.

### Experiment I: artificial selection experiment

Details of the experimental procedure for artificial selection can be found in Zu & Schiestl [14]. To summarize, we sowed out 150 seeds to form the parental generation. Up and down directional artificial selection on plant height were imposed to produce a tall and a short line with the ten tallest and ten shortest plants, respectively. Additionally, ten randomly selected plants were chosen to form a control line. Selected plants were randomly hand pollinated within each line. Pollen donor, pollen receiver and their offspring were labeled for each fruit to generate a breeding pedigree. After fruit maturation, around 50 seeds from each of the three lines were sown out to form the next generation. The same procedures were carried out to obtain three generations of selection. Extra seeds were sowed out to ensure a minimum of 150 individual plants in each generation. In total, we analyzed 628 plants. The experiment was conducted in a phytotron with 24h fluorescent light per day, 22°C, 60% relative humidity, and regular watering twice a day (at 08:00 and 18:00).

### Experiment II: pollinator selection experiments

The procedures of experimental evolution experiment can be found in detail in Gervasi & Schiestl [25]. To summarize, we sowed out 300 seeds to generate 108 full sib families by manual cross pollination. These 108 full sib families were then equally divided into three replicates each containing 36 plants, for each of the three treatments (bumblebee, hoverfly, and hand pollination treatment). In each replicate, the 36 plants were placed in a 6*6 array with a distance of 20 cm from each other in a flight cage (2.5m*1.8m*1.2m). In bumblebee and hoverfly treatments, five pollinators (either *Bombus terrestris* or *Episyrphus balteatus*) were introduced one at a time in the flight cage, with each allowed to freely visit maximal three different plants before being removed from the cage. A total number of 12 – 15 out of 36 plants per replicate received one or more pollinator visitation. The average (± s.d.) visitation (in visited plants) was 1.35 ± 0.63 for bumblebee-pollinated plants and 1.28 ± 0.53 for hoverfly-pollinated plants. In the control treatment 12 plants were randomly chosen and were manually pollinated among each other. Floral traits were measured prior to pollinators’ visits or hand pollination. The number of seeds were recorded after fruit maturation. Seeds from the pollinated plants were sown out proportionally (36/(replicate sum of seeds/individual seed set), values below 0.5 were rounded up to 1) to form again a total number of 36 plants for the next generation of each replicate. The same selection and sowing-out procedures were conducted for 9 generations, after which plants were sowed out again and randomly hand crossed between the replicates within each treatment to get rid of potential inbreeding depression. Fruits from random crosses were sown out to form the 11^th^ generation and the measurements of floral traits in this generation were used as observed responses to selection.

### Estimation of the genetic variance-covariance matrices (G-matrix)

With known breeding pedigree and plant trait values for each individual in the control and treatment lines of the artificial selection experiment, we were able to estimate three genetic variance-covariance matrices: **G**_control_ in control, **G**_tall_ (or **G**_short_) in selection lines for increased (or decreased) plant height (see Table S3). The pedigree of the seeds sowed in the pollinator experiment was unknown. We thus used **G**_control_ from artificial selection experiment for evolutionary predictions in both experiments. More specifically, we estimated the G-matrix of the 16 traits by using a multivariate animal model in which the kinship (relatedness) matrix was obtained from the four-generation pedigree of the plants crossed within the experiments (sire = pollen donor, dam = pollen receiver), independently in the control, tall, and short experimental lines (Table S3). We fitted a linear mixed model using the Bayesian method implemented in the MCMCglmm R package [63] to estimate random effect variance components for additive genetic effects (V_A_) from which we estimated the G-matrix, and among-dam (V_D_) and among-sire (V_S_) components to remove potential maternal and paternal effects, respectively. We added generation as a block factor modeled as a fixed effect. This method was previously shown to have good applications with a few traits [48, 64].

In MCMCglmm, we used weakly informative inverse-Wishart prior with limit variance of one and covariance of zero and low degree of belief (0.002). Posterior distributions were robust to several different prior settings (e.g. V = diag(n)*0.1, V = diag(n)*10, n = number of traits).

We used 1,200,000 iterations, with a burn-in of 200,000 and a thinning of 500 to ensure convergence and low autocorrelation among thinned samples (< 0.1). The thinning resulted in a posterior distribution with 2000 samples.

Finally, because the Bayesian approach does not allow us to directly test for the accuracy of our estimates of the G-matrices, we implemented a permutation test in which we randomly shuffled the dam and sire of each offspring within each generation and re-estimated the G-matrix for each of 500 replicates using the same MCMCglmm procedure as before. To evaluate the accuracy of the observed G-matrices (**G**_control_, **G**_tall_, and **G**_short_), we then compared them to their randomized estimates, element by element. For each element, we computed an empirical *P*-value as: *P = (N_random.estimates_ < observed.value)/500*. If the observed value was smaller than the mean of the random estimates, then (1 − *P*) was used instead of *P*. The random estimates were obtained from the posterior mode of the 500 random estimates of each G-matrix. An element of **G** (a variance or covariance term within **G**) was considered significant if its *P*-value was < 0.05. If it was not the case, then the specific element estimation did not capture its biological meaning.

### Estimation of selection gradients

In the artificial selection experiment, we calculated the selection gradient on height (*β_h_*) by using

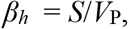

where *V*_p_ is the phenotypic variation of height and *S* the selection differential calculated as the difference between the mean plant height of the selected plants and all measured plants in the same generation.

We calculated *β_h_* in each generation and each selected line (Table S1) and used its sum over the three generations to predict the total evolutionary responses in each line.

In the pollinator selection experiments, we estimated the selection gradients following the partial correlation approach of Lande and Arnold [31]. We used generalized linear models with fitness as dependent variable and morphological and scent variable as covariates. We estimated fitness as the total number of seeds produced per plant (female fitness component) plus a male fitness component gained through pollen export. We calculated the male component as the product of “number of pollinator visits” (i.e. number of visits by pollinators to a given plant) times their respective efficiency. Pollinator efficiency is the average of the resulting number of “seeds per fruit” per single pollinator visit to a plant across all generations. Pollinator efficiency was strongly species dependent (mean ± S.D.: bumblebee: 10.57±6.34; hoverfly: 4.81±3.83; t_267_=10.11, P<0.001). Fitness was ln(1+x) transformed before analysis to reach normality. The average selection gradients (**β**) were calculated separately per treatment (bumblebee, hoverfly, and control), for all the measured generations and replicates combined (for details, see [25]). Therefore, we used 9\***β** to predict the total evolutionary changes after 9 generations of pollinator selection. The non-significant selection gradients were still used as the best approximate estimations of selection.

### Calculation of predicted and observed evolutionary changes

To estimate the predicted responses to selection, we used the multivariate breeder’s equation [30], Δ**z** = **G*β** (see Introduction). We used **G** from the control group in the artificial selection experiment (**G**_control_, Table S3, S4) for predictions as the best estimation of genetic architecture of the original population. We used the 2000 posterior samples of the G-matrix to generate a distribution of predicted trait changes from which we could evaluate the accuracy of our evolutionary prediction using its 95% highest posterior density (HPD) interval.

To calculate the observed trait changes, we did three step calculations as follows (Fig. S1). 1) We calculated the absolute phenotypic changes between the last and the first generation (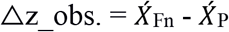, where n is 3 in artificial selection experiment, and 11 in pollinator selection experiment) for each line or each treatment, and P stands for ‘parental’. 2) We estimated environmental fluctuations by using the control groups. Ideally, there should be no selection and no systematic changes of trait values in the control groups beside random fluctuations. However, traits values may have changed due to unaccounted-for selection pressures unknown to the experimenters. Therefore, we subtracted the predicted changes due to unknown selection (**G* β_control_** = Δ**z**__control_pred._) from the observed changes in control group to estimate environmental fluctuations. We calculated environmental fluctuations in both artificial- and pollinator-selection experiments. 3) We subtracted the environmental fluctuations (values in step 2) from the absolute changes (values in step 1) and used the resulting values as the observed evolutionary changes to compare with predicted changes. We present the observed and predicted changes scaled by the phenotypic standard deviation of each trait in the parental generation.

### Direct and indirect selection responses

To examine the importance of trait covariance in affecting evolutionary trajectories, we separated the total selection response Δ**z** of each trait into its direct and indirect components. The direct component of the predicted selection response of trait *i* is the product G*_ii_*\*β*_i_*, with G*_ii_* the additive genetic variance of the trait (diagonal element of **G**_control_). The indirect component is the product of the off-diagonal elements of **G** (genetic covariance: G*_ij_*) with **β**, summed over all traits *j* ≠ *i*: 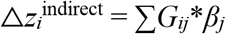. The total response is the sum of these two components. The three predictions, indirect, direct, and total response were compared to the observed change of each trait to evaluate when the direct response is constrained (direct and indirect components of opposite sign) or enhanced (direct and indirect components of same sign) by genetic covariance.

Finally, we measured the constraining effect of genetic co-variation on the response to selection by comparing the angle **θ** between the selection response vector (Δ**z**) and the first PC of **G** (PC1, or ***g***_max_) with the angle **γ** between Δ**z** and the selection gradient (β). We generated the posterior distribution of **θ** from the posterior distribution of **G**_control_, which allowed us to test whether **γ** is larger (smaller) than **θ**, which tests if Δ**z** is biased (unbiased) in the direction of ***g***_max_ by genetic correlations.

### G-matrices similarity among artificial selection lines

We compared the **G**-matrices from control, tall and short selection lines to assess the stability of **G** between treatments and control. We used the random skewers (RS) method in one comparison test because it examines the similarity between two **G**-matrices of their expected evolutionary response to a random set of selection vectors (skewers), which fits our purpose of evaluating the stability of such predictions using different estimates of the **G**-matrix. We used Roff et al.’s (2012) implementation of the RS method, and report the mean over 10,000 random selection skewers of the correlation between the selection response vectors of the two G-matrices compared. Significance was obtained from the distribution of the test statistics obtained from the 500 random estimates of each **G**-matrix. We performed a further test of shape similarity between the **G**-matrices using the hierarchical approach of Roff et al. [65], also known as the Flury hierarchy [66]. This method tests the degree of shape similarity sequentially by comparing the size and orientation of the eigenvectors (principal components, PCs) of the **G**-matrices. Two **G**-matrices can have common principal components (CPC) if their PCs have the same orientation but not the same size (i.e., have different eigenvalues), be proportional if their PCs only differ proportionally, or be equal. The three levels of similarity are tested relative to the hypothesis of unrelated matrices. The test statistics are provided in Roff et al. [65]. We determined the significance of the RS and Flury tests using the previous 500 randomized estimates of **G**_control_, **G**_tall_, and **G**_short_.

All statistics were conducted with R version 3.3.3. [67].

## Supporting information

Details of some methods parts, *i.e*. floral scent collection and analysis, and comparing shape, size and orientation of **G**-matrices; as well as some additional supporting result tables and figures that cited in the main text can be found in the Supporting information part of this article.

## Declarations

### Ethics approval and consent to participate

Not applicable.

### Consent for publication

Not applicable.

### Availability of data and materials

The datasets generated and/or analysed during the current study are available in the Dryad repository, [PERSISTENT WEB LINK TO DATASETS].

### Authors’ contributions

PZ, FS, DG performed experimental data analysis; PZ, FG performed most of the evolutionary statistical analysis; XL, DR performed part of the prediction analysis; PZ, FS, FG wrote the manuscript; all authors revised the manuscript.

## Acknowledgements

We thank Jarrod Hadfield for his fast and helpful reply whenever we had questions related to MCMCglmm approach.

