## Supplementary for "Floral signals evolve in a predictable way under artificial and pollinator selection in *Brassica rapa* using a G-matrix"

### Supporting Information

#### Material and methods

##### *Scent collection and analysis*

The measurement methods were described in detail in Zu & Schiestl (2017). We collected floral volatiles of the entire inflorescence, when at least four freshly opened flowers were present, by using a push-pull system, and analyzed floral VOCs by using a gas chromatograph with a mass-selective detector (GC-MSD) fitted with a thermal desorption system (Gerstel TDS/TDU, Mülheim an der Ruhr, Germany). With the push-pull scent collection system, the whole inflorescence was fixed with two Teflon plates (with an opening in the middle for the stem) and was covered in a glass cylinder with silanized (deactivated) surfaces (Sigmacote, Sigma, MO, USA). Two glass ports were used to push-in clean air and pull-out air with volatiles for sampling. Through the push port we introduced a filter with activated charcoal (SKC Eighty Four, PA, USA) which was connected to an air pump to push purified air at 100 mL per min into the glass cylinder. Through the pull port we introduced a glass tube filled with adsorbent (35 mg Tenax TA 60/80, Supelco, Bellefonte, PA, USA) which was connected to a vacuum pump for pulling out air containing inflorescence VOCs at a flow rate of 100 mL per min, for 3 hours for each plant. Wisconsin's rapid cycling *B. rapa* has been bred under 24 hours light a day. No significant daily dynamics of scent emission were found in a pilot experiment (scent collection for 3 plants during three different time periods of a day, data not shown). Thus, we collected scent during the following time periods: 07:00 – 10:00, 12:00 – 15:00, and 17:00 – 20:00 hours, depending on the number of available plants. At least one air control from an empty glass cylinder was sampled in each collection period.

To analyze floral VOCs, each sample was injected into a gas chromatograph (GC, Agilent 6890N, Agilent Technologies, Palo Alto, CA, USA) using a Gerstel Multipurpose Sampler. For thermal desorption, a temperature program was run, starting at 30°C (1 minute hold) increasing to 240°C (1 minute hold) at 60°C min$^{-1}$. A cool injection system (CIS 4, Gerstel Germany) was used for cryotrap enrichment of eluting volatiles from the TDS. The CIS started with a temperature of -150°C (0.05 minute hold) and increased to 150°C at 16°C s$^{-1}$, then increased further from 150°C to 250°C at 12°C s$^{-1}$. The GC was equipped with an HP-5 capillary column (Agilent, 15m length, 0.25mm diameter, 0.25µm film thickness). The temperature of the GC oven was set to 50°C (1 minute hold) and increased to 250°C at 10°C min$^{-1}$, and helium was used as the carrier gas, with a constant flow of 2 ml min$^{-1}$. An Agilent

5975 Series MSD mass spectrometer was used for compound identification and quantification.

We used an Agilent 5975 Series MSD mass spectrometer for compound identification and quantification. To identify volatile compounds, mass spectra obtained from the samples were compared with those of a reference collection (NIST, the National Institute of Standards and Technology, mass spectral library). Subsequently, retention times and mass spectra of all compounds included in the quantitative analyses were compared with those of synthetic reference standards. Compound quantification was carried out with the ChemStation Enhanced Data Analysis program (Version E.01.00). For all compounds included in the study, synthetic standards were analyzed in three different amounts using the GC-MSD (1, 10 and 100 ng). By using target ions for each compound, calibration curves were established using the ChemStation program. Peak areas of target ions were subsequently used for quantifying the amounts of volatiles in the samples.

***Comparing shape, size and orientation of G-matrices***

Besides the random skewer method to compare the similarity of G-matrices, we further evaluated the size, the shape, and the orientation of the G-matrices (Puentes *et al.*, 2016) in the control-, tall-, and short-lines by examining their posterior G-matrices obtained from *MCMCglmm* models.

First, the shape of the G-matrices were compared by using a principle component (PC) analysis to characterize the main axes and the loadings of the traits on the PCs. We calculated the eigenvectors and eigenvalues ($\lambda_i$, i=1, 2, 3,…, p) by performing singular value decomposition (SVD) of G-matrix(Golub & van Loan, 1983) using the *eigen* function in R v3.3.3. Because of the orthogonality of the eigenvectors, we can estimate the shape of a matrix by comparing their associated eigenvalues $\lambda_i$ and their contribution to the total genetic variation of the traits:

$$\frac{\lambda_i}{\sum_{i=1}^{p} \lambda_i}.$$

Similar eigenvalues in different axes (eigenvectors) indicate a more ‘spherical’ shape in multivariate trait space, whereas dissimilar eigenvalues indicate a more ‘elliptical’ shape.

Second, the size of a G-matrix, corresponding to the total genetic variance of the set of traits, sometimes referred as the volume of the G-matrix, is estimated as the sum of the eigenvalues:

$\sum_{i=1}^{p} \lambda_i$.

The bigger the size is, the larger the total amount of genetic variation in the set of traits in the population.

Third, we used the orientation of the first important PC (PC1) as the representative for the major orientation of a G-matrix ($\boldsymbol{g}_{max}$). We examined the angles of PC1s within and between posterior distributions of the four **G**s by using the equation of angles between two non-zero vector *u* and *v*: $cos\theta = \frac{u \cdot v}{\|u\| \|v\|}$.

**Table S1**. Selection gradients on plant height in *Brassica rapa* in the tall and short artificial selection experiment for three generations.

| | **Tall** | **Short** |
|---|---|---|
| **β1** | 0.3543 | -0.2828 |
| **β2** | 0.3313 | -0.2153 |
| **β3** | 0.2914 | -0.3872 |
| **Total** | 0.9770 | -0.8853 |

**Table S2**. Linear and quadratic selection gradients and standard error on various traits in *Brassica rapa* in bumblebee, hoverfly selection and control groups. We used pooled data from all the generations and all replicates in each experiment. Significant values are bolded.

| | Bumblebee | | | | Hoverfly | | | | Control | | | |
|---|---|---|---|---|---|---|---|---|---|---|---|---|
| | Linear selection | | Quadratic selection | | Linear selection | | Quadratic selection | | Linear selection | | Quadratic selection | |
| **Traits** | β | SE | γ | SE | β | SE | γ | SE | β | SE | γ | SE |
| **Plant height** | **0.487** | 0.103 | 0.107 | 0.060 | 0.114 | 0.079 | -0.068 | 0.051 | 0.035 | 0.102 | -0.033 | 0.071 |
| **Petal width** | 0.098 | 0.113 | 0.025 | 0.071 | -0.101 | 0.081 | -0.004 | 0.053 | -0.046 | 0.109 | -0.070 | 0.073 |
| **Petal length** | 0.243 | 0.141 | -0.120 | 0.081 | 0.054 | 0.106 | -0.063 | 0.065 | 0.251 | 0.131 | -0.068 | 0.074 |
| **Flower diameter** | -0.069 | 0.141 | 0.135 | 0.070 | 0.148 | 0.105 | 0.096 | 0.066 | 0.010 | 0.130 | 0.047 | 0.072 |
| **Benzaldehyde** | **-0.301** | 0.135 | -0.109 | 0.086 | -0.130 | 0.112 | -0.122 | 0.066 | -0.098 | 0.123 | -0.035 | 0.069 |
| **Phenylacetaldehyde** | **0.441** | 0.189 | **0.235** | 0.114 | -0.148 | 0.139 | 0.149 | 0.098 | 0.015 | 0.177 | 0.085 | 0.108 |
| **Phenylethyl alcohol** | -0.135 | 0.172 | -0.064 | 0.075 | -0.016 | 0.125 | -0.039 | 0.077 | 0.004 | 0.158 | -0.071 | 0.070 |
| **Methyl salicylate** | -0.009 | 0.146 | 0.074 | 0.100 | 0.084 | 0.105 | -0.083 | 0.066 | -0.114 | 0.123 | **0.192** | 0.082 |
| **Methyl benzoate** | **0.505** | 0.161 | -0.067 | 0.095 | -0.080 | 0.127 | 0.029 | 0.081 | 0.142 | 0.132 | -0.056 | 0.070 |
| **Methyl anthranilate** | -0.180 | 0.188 | -0.101 | 0.105 | -0.263 | 0.137 | -0.133 | 0.074 | -0.012 | 0.153 | 0.045 | 0.090 |
| **Benzyl nitrile** | **-0.557** | 0.209 | -0.037 | 0.123 | 0.169 | 0.134 | 0.138 | 0.080 | -0.111 | 0.194 | 0.161 | 0.114 |
| **2-Amino benzaldehyde** | 0.162 | 0.176 | 0.064 | 0.110 | -0.027 | 0.121 | 0.044 | 0.079 | 0.265 | 0.170 | -0.036 | 0.108 |
| **Indole** | **0.420** | 0.194 | -0.182 | 0.126 | 0.176 | 0.136 | **-0.236** | 0.086 | -0.069 | 0.171 | -0.126 | 0.097 |
| **(E)-α-Farnesene** | -0.101 | 0.128 | -0.024 | 0.068 | 0.186 | 0.100 | 0.106 | 0.064 | **0.338** | 0.115 | 0.015 | 0.069 |
| **Z-(3)-Hexenyl acetate** | -0.080 | 0.121 | -0.015 | 0.090 | -0.174 | 0.103 | -0.020 | 0.056 | -0.029 | 0.113 | -0.148 | 0.088 |
| **1-Butene-4-isothyocy.** | -0.189 | 0.114 | 0.022 | 0.085 | -0.004 | 0.095 | -0.103 | 0.065 | 0.116 | 0.108 | -0.009 | 0.074 |

**Table S3.** Different model setups for MCMCglmm analyses of G-matrices estimation. Models include the three models from different lines throughout the artificial selection experiment in *Brassica rapa*. Generation 0, parental generation; 1, 2, 3, offspring generations. N, plant sample size for each model.

| **G names** | **Line** | **Generation** | **Fixed factor** | **Random factor** | **N** |
|---|---|---|---|---|---|
| $G_{control}$ | Control | 0,1,2,3 | Generation | Dam, sire | 298 |
| $G_{tall}$ | Tall | 0,1,2,3 | Generation | Dam, sire | 298 |
| $G_{short}$ | Short | 0,1,2,3 | Generation | Dam, sire | 290 |

**Table S4.** Posterior mode of genetic variance (diagonal), covariance (lower triangular) and genetic correlations (upper triangular) estimated from control, tall, and short lines in the artificial selection experiment. Significant values are bolded (uncorrected values).

| | **Height** | **PW** | **PL** | **FD** | **Ben** | **PAA** | **FAR** | **BenN** | **Aben** | **Ind** | **MA** | **PA** | **MS** | **MB** | **ZHA** | **ITC** |
|---|---|---|---|---|---|---|---|---|---|---|---|---|---|---|---|---|
| ***Control*** | | | | | | | | | | | | | | | | |
| Height | **23.192** | **0.142** | **0.272** | **0.437** | **0.165** | **0.275** | **0.282** | **0.113** | **0.285** | **0.322** | **0.311** | **0.296** | **0.054** | **0.184** | **0.190** | **0.139** |
| PW | **0.369** | **0.292** | **0.292** | **0.380** | **0.051** | **0.155** | 0.082 | **0.116** | **0.141** | **0.122** | **0.154** | **0.173** | 0.076 | **0.088** | **0.020** | -0.017 |
| PL | **0.704** | **0.085** | **0.288** | **0.521** | **0.030** | **0.171** | **0.163** | **0.131** | **0.166** | **0.190** | **0.230** | **0.142** | 0.034 | **0.139** | **0.045** | 0.031 |
| FD | **1.978** | **0.193** | **0.263** | **0.881** | **0.033** | **0.245** | **0.165** | **0.156** | **0.245** | **0.254** | **0.294** | **0.286** | **0.088** | **0.167** | **0.080** | **0.117** |
| Ben | **0.493** | **0.017** | **0.010** | **0.019** | 0.384 | **0.163** | **0.152** | **0.141** | **0.228** | **0.176** | **0.103** | **0.069** | 0.057 | **0.204** | 0.117 | **0.053** |
| PAA | **1.297** | **0.082** | **0.090** | **0.226** | **0.099** | **0.963** | **0.244** | **0.360** | **0.412** | **0.324** | **0.329** | **0.664** | **0.243** | **0.202** | **0.132** | **0.190** |
| FAR | **0.744** | 0.024 | **0.048** | **0.085** | **0.052** | **0.131** | **0.300** | **0.196** | **0.295** | **0.246** | **0.262** | **0.273** | 0.197 | **0.149** | 0.141 | **0.192** |
| BenN | **0.328** | **0.038** | **0.042** | **0.088** | **0.053** | **0.213** | **0.065** | **0.364** | **0.325** | **0.368** | **0.269** | **0.428** | 0.100 | **0.200** | 0.047 | **0.115** |
| Aben | **1.264** | **0.070** | **0.082** | **0.212** | **0.130** | **0.372** | **0.149** | **0.181** | **0.848** | **0.631** | **0.481** | **0.324** | **0.152** | **0.156** | **0.151** | **0.160** |
| Ind | **1.100** | **0.047** | **0.072** | **0.169** | **0.077** | **0.225** | **0.096** | **0.157** | **0.412** | **0.502** | **0.399** | **0.363** | **0.234** | **0.139** | **0.181** | **0.205** |
| MA | **1.215** | **0.067** | **0.100** | **0.224** | **0.052** | **0.262** | **0.116** | **0.132** | **0.360** | **0.229** | **0.658** | **0.348** | **0.300** | **0.429** | 0.117 | 0.096 |
| PA | **1.077** | **0.070** | **0.057** | **0.202** | **0.032** | **0.492** | **0.113** | **0.195** | **0.225** | **0.194** | **0.213** | **0.569** | **0.253** | **0.266** | **0.190** | **0.289** |
| MS | **0.197** | 0.032 | 0.014 | **0.063** | 0.027 | **0.182** | 0.082 | 0.046 | **0.106** | **0.126** | **0.185** | **0.145** | **0.581** | **0.314** | 0.124 | 0.035 |
| MB | **0.624** | **0.034** | **0.052** | **0.111** | **0.089** | **0.140** | **0.058** | **0.085** | **0.101** | **0.069** | **0.246** | **0.142** | **0.169** | **0.498** | **0.181** | **0.199** |
| ZHA | **0.586** | **0.007** | **0.015** | **0.048** | 0.047 | **0.083** | 0.049 | 0.018 | **0.089** | **0.082** | 0.061 | **0.092** | 0.061 | **0.082** | **0.412** | **0.169** |
| ITC | **0.579** | -0.008 | 0.014 | **0.095** | **0.028** | **0.162** | **0.091** | **0.060** | **0.128** | **0.126** | 0.067 | **0.190** | 0.023 | **0.122** | **0.094** | 0.753 |
| ***Tall*** | | | | | | | | | | | | | | | | |
| Height | **2.075** | 0.021 | **0.171** | **0.272** | 0.086 | 0.027 | 0.039 | **-0.106** | 0.018 | 0.068 | 0.130 | **0.119** | -0.058 | **0.121** | 0.069 | -0.040 |
| PW | 0.016 | 0.293 | **0.291** | 0.334 | **0.111** | 0.073 | 0.071 | **0.139** | 0.083 | 0.021 | **0.125** | **0.101** | 0.075 | **0.092** | **0.030** | -0.103 |
| PL | **0.132** | **0.085** | **0.287** | **0.546** | 0.053 | 0.091 | **0.056** | **0.143** | **0.120** | 0.091 | 0.110 | 0.079 | **0.071** | **0.109** | -0.031 | -0.006 |
| FD | **0.360** | 0.166 | **0.269** | **0.846** | **0.057** | **0.221** | **0.106** | **0.133** | 0.070 | 0.078 | 0.101 | **0.138** | **0.101** | **0.091** | **0.038** | -0.090 |
| Ben | 0.076 | **0.037** | 0.017 | **0.032** | **0.378** | 0.054 | 0.150 | 0.112 | **0.227** | **0.150** | **0.103** | **0.129** | **0.117** | **0.296** | **0.222** | 0.026 |
| PAA | 0.033 | 0.034 | 0.042 | **0.173** | 0.028 | **0.726** | **0.190** | **0.341** | **0.312** | **0.238** | **0.268** | **0.574** | **0.117** | 0.051 | 0.010 | 0.079 |
| FAR | 0.030 | 0.021 | **0.016** | **0.053** | 0.050 | **0.088** | **0.295** | 0.149 | **0.235** | **0.244** | 0.159 | **0.202** | **0.220** | **0.138** | **0.144** | 0.136 |
| BenN | **-0.089** | **0.044** | **0.044** | **0.071** | 0.040 | **0.169** | 0.047 | **0.337** | **0.338** | **0.337** | **0.206** | **0.289** | **0.188** | **0.115** | **0.129** | 0.009 |
| Aben | 0.020 | 0.035 | **0.050** | 0.051 | **0.110** | **0.208** | **0.100** | **0.154** | 0.615 | **0.523** | 0.439 | **0.263** | **0.182** | **0.100** | 0.103 | 0.103 |
| Ind | 0.066 | 0.008 | 0.033 | 0.048 | **0.062** | **0.137** | **0.089** | **0.132** | **0.276** | **0.453** | **0.370** | **0.170** | **0.152** | **0.049** | 0.093 | 0.109 |
| MA | 0.139 | **0.050** | 0.044 | 0.069 | **0.047** | **0.170** | 0.064 | **0.089** | 0.256 | **0.185** | **0.553** | **0.211** | **0.274** | **0.336** | **0.161** | **0.162** |
| PA | **0.115** | **0.037** | 0.029 | **0.086** | **0.053** | **0.329** | **0.074** | **0.113** | **0.139** | **0.077** | **0.106** | **0.452** | **0.125** | **0.200** | 0.128 | **0.205** |

| | | | | | | | | | | | | | | | | |
|---|---|---|---|---|---|---|---|---|---|---|---|---|---|---|---|---|
| MS | -0.063 | 0.030 | **0.028** | **0.069** | **0.053** | **0.074** | **0.089** | **0.081** | **0.106** | **0.076** | **0.152** | **0.063** | **0.553** | **0.358** | 0.063 | **0.050** |
| MB | **0.120** | **0.034** | **0.040** | **0.058** | **0.125** | 0.030 | **0.052** | **0.046** | **0.054** | **0.023** | **0.172** | **0.093** | **0.183** | **0.474** | 0.200 | 0.074 |
| ZHA | 0.058 | **0.009** | -0.010 | **0.020** | **0.079** | 0.005 | **0.045** | **0.043** | 0.047 | 0.037 | **0.069** | 0.050 | 0.027 | 0.080 | 0.337 | **0.133** |
| ITC | -0.047 | -0.045 | -0.003 | -0.068 | 0.013 | 0.055 | 0.061 | 0.004 | 0.066 | 0.060 | **0.099** | **0.113** | **0.030** | 0.042 | **0.063** | 0.669 |
| ***Short*** | | | | | | | | | | | | | | | | |
| Height | 23.163 | 0.119 | 0.247 | 0.254 | **0.370** | **0.235** | 0.222 | **0.216** | **0.437** | **0.448** | **0.387** | **0.194** | 0.011 | **0.306** | 0.274 | **0.118** |
| PW | 0.326 | **0.323** | **0.282** | **0.335** | **0.092** | **0.140** | 0.034 | **0.136** | **0.097** | **0.114** | **0.168** | **0.091** | 0.056 | **0.125** | 0.031 | **-0.008** |
| PL | 0.649 | **0.087** | **0.298** | **0.560** | **0.082** | **0.196** | **0.078** | 0.094 | **0.158** | **0.166** | **0.166** | **0.068** | 0.019 | **0.085** | **0.064** | -0.029 |
| FD | 1.098 | **0.171** | **0.275** | **0.809** | 0.045 | **0.214** | **0.087** | **0.120** | **0.258** | **0.189** | **0.241** | **0.221** | -0.014 | **0.129** | **0.102** | **0.018** |
| Ben | **1.149** | **0.034** | **0.029** | 0.026 | **0.417** | **0.155** | **0.210** | 0.076 | **0.289** | 0.193 | **0.188** | 0.096 | **0.176** | **0.360** | **0.245** | 0.070 |
| PAA | **1.048** | **0.074** | **0.099** | **0.178** | **0.093** | **0.861** | **0.227** | 0.311 | **0.276** | **0.308** | **0.302** | **0.609** | 0.123 | **0.231** | **0.148** | **0.145** |
| FAR | 0.822 | 0.015 | **0.033** | **0.060** | **0.104** | **0.162** | **0.592** | 0.102 | **0.210** | **0.240** | **0.176** | 0.152 | **0.154** | 0.067 | **0.145** | 0.119 |
| BenN | **0.603** | **0.045** | 0.030 | **0.063** | 0.028 | 0.167 | 0.046 | **0.335** | **0.330** | **0.303** | **0.268** | 0.248 | 0.149 | **0.169** | **0.117** | 0.060 |
| Aben | **1.783** | **0.047** | **0.073** | **0.196** | **0.158** | **0.217** | **0.137** | **0.162** | **0.718** | **0.539** | **0.565** | **0.246** | 0.130 | **0.194** | **0.171** | **0.140** |
| Ind | **1.468** | **0.044** | **0.062** | **0.116** | 0.085 | **0.195** | **0.126** | **0.120** | **0.311** | **0.464** | **0.428** | **0.255** | 0.162 | **0.191** | **0.223** | **0.138** |
| MA | **1.509** | **0.077** | **0.073** | **0.176** | **0.099** | **0.227** | **0.110** | **0.126** | **0.388** | **0.237** | **0.658** | **0.238** | **0.201** | **0.430** | **0.332** | **0.129** |
| PA | **0.666** | **0.037** | **0.026** | **0.142** | 0.044 | **0.404** | 0.083 | 0.103 | **0.149** | **0.124** | **0.138** | **0.511** | 0.108 | **0.255** | **0.170** | 0.088 |
| MS | 0.039 | 0.023 | 0.008 | -0.009 | **0.085** | 0.085 | **0.088** | 0.064 | 0.082 | **0.082** | **0.121** | 0.058 | **0.553** | 0.259 | 0.120 | **0.090** |
| MB | **0.992** | **0.048** | **0.031** | **0.078** | **0.156** | **0.144** | 0.035 | **0.066** | **0.111** | **0.088** | **0.235** | **0.123** | 0.130 | **0.452** | **0.302** | 0.131 |
| ZHA | 0.933 | 0.012 | **0.025** | **0.065** | **0.112** | **0.097** | **0.079** | **0.048** | **0.102** | **0.108** | **0.191** | **0.086** | 0.063 | **0.144** | **0.501** | **0.221** |
| ITC | **0.528** | **-0.004** | -0.015 | **0.015** | 0.042 | **0.125** | 0.085 | 0.032 | **0.110** | **0.088** | **0.097** | 0.058 | **0.062** | 0.082 | **0.145** | **0.865** |

**Table S5.** Eigen-structure (eigenvalues and PC1) of posterior G-matrices.

| | Eigenvalues | | | PC1 | | |
|---|---|---|---|---|---|---|
| | $\mathbf{G}_{control}$ | $\mathbf{G}_{tall}$ | $\mathbf{G}_{short}$ | $\mathbf{G}_{control}$ | $\mathbf{G}_{tall}$ | $\mathbf{G}_{short}$ |
| Height | 23.7743 | 2.7994 | 22.3011 | 0.9863 | 0.9687 | 0.9835 |
| PW | 2.1650 | 1.4876 | 1.7313 | 0.0235 | 0.0447 | 0.0205 |
| PL | 1.0032 | 0.9289 | 1.0776 | 0.0300 | 0.0581 | 0.0290 |
| FD | 0.7969 | 0.7957 | 0.7964 | 0.0881 | 0.1254 | 0.0616 |
| Ben | 0.6700 | 0.6059 | 0.6578 | 0.0250 | 0.0400 | 0.0439 |
| PAA | 0.5268 | 0.5207 | 0.5439 | 0.0640 | 0.0612 | 0.0614 |
| FAR | 0.4185 | 0.4199 | 0.4438 | 0.0381 | 0.0306 | 0.0442 |
| BenN | 0.3622 | 0.3452 | 0.3798 | 0.0154 | 0.0005 | 0.0217 |
| Aben | 0.3142 | 0.3101 | 0.3415 | 0.0705 | 0.1147 | 0.0888 |
| Ind | 0.2613 | 0.2570 | 0.2910 | 0.0464 | 0.0388 | 0.0659 |
| MA | 0.2444 | 0.2326 | 0.2548 | 0.0539 | 0.1161 | 0.0776 |
| PA | 0.2133 | 0.2041 | 0.2260 | 0.0493 | 0.0749 | 0.0291 |
| MS | 0.1889 | 0.1793 | 0.2048 | 0.0162 | 0.0384 | 0.0235 |
| MB | 0.1698 | 0.1663 | 0.1783 | 0.0355 | 0.0708 | 0.0512 |
| ZHA | 0.1518 | 0.1480 | 0.1559 | 0.0279 | 0.0112 | 0.0476 |
| ITC | 0.1279 | 0.1263 | 0.1319 | 0.0317 | -0.0478 | 0.0427 |

**Table S6:** Observed and predicted trait changes in the (A) Bumblebee and (B) Hoverfly experiments. The predicted changes are obtained from the multivariate breeder's equation and are decomposed into the direct and indirect components of the total response to selection (see main text). Figures in bold are predicted responses who's 95% HPD interval does not overlap with zero or observed changes who's value falls within the 95% HPD interval of its total predicted value.

| | Observed changes | Predicted changes | | | | | | | | |
|---|---|---|---|---|---|---|---|---|---|---|
| | | Total | 95% HPD | | Direct | 95% HPD | | Indirect | 95% HPD | |
| | | | lower | upper | | lower | upper | | lower | upper |
| ***(A) Bumblebee*** | | | | | | | | | | |
| Plant height (Height) | 1.361 | **2.888** | 1.989 | 3.637 | **2.660** | 1.782 | 3.370 | **0.188** | 0.015 | 0.364 |
| Petal width (PW) | **0.267** | **0.635** | 0.090 | 1.077 | **0.050** | 0.038 | 0.066 | **0.584** | 0.043 | 1.024 |
| Petal length (PL) | **0.423** | **0.893** | 0.300 | 1.403 | **0.135** | 0.101 | 0.178 | **0.753** | 0.156 | 1.228 |
| Flower diameter (FD) | 0.281 | **0.690** | 0.377 | 1.206 | **-0.044** | -0.064 | -0.032 | **0.853** | 0.429 | 1.280 |
| Benzaldehyde (Ben) | 0.375 | 0.237 | -0.463 | 1.108 | **-0.249** | -0.337 | -0.188 | 0.549 | -0.194 | 1.403 |
| Phenylacetaldehyde (PAA) | 1.746 | **0.690** | 0.247 | 1.404 | **0.356** | 0.236 | 0.551 | 0.373 | -0.136 | 0.904 |
| α-Farnesene (FAR) | **0.859** | **0.650** | 0.208 | 1.077 | **-0.041** | -0.058 | -0.033 | **0.697** | 0.256 | 1.136 |
| Benzyl nitrile (BenN) | 1.744 | 0.032 | -0.302 | 0.563 | **-0.231** | -0.305 | -0.164 | 0.379 | -0.052 | 0.862 |
| 2-Amino benzaldehyde (Aben) | 1.220 | **0.565** | 0.119 | 0.977 | **0.092** | 0.051 | 0.120 | **0.470** | 0.053 | 0.893 |
| Indole (Ind) | **1.200** | **0.647** | 0.292 | 1.230 | **0.201** | 0.140 | 0.282 | **0.542** | 0.119 | 0.984 |
| Methyl anthranilate (MeA) | **1.158** | **0.540** | 0.100 | 1.243 | **-0.118** | -0.168 | -0.075 | **0.834** | 0.212 | 1.389 |
| Phenylethyl alcohol (PA) | 2.809 | **1.283** | 0.229 | 2.465 | **-0.161** | -0.260 | -0.122 | **1.440** | 0.417 | 2.708 |
| Methyl salicylate (MeS) | -0.437 | 0.295 | -0.340 | 0.829 | **-0.006** | -0.009 | -0.004 | 0.303 | -0.334 | 0.836 |
| Methyl benzoate (MeB) | **1.011** | **0.805** | 0.196 | 1.340 | **0.330** | 0.235 | 0.443 | 0.398 | -0.111 | 0.942 |
| Z-(3)-Hexenyl acetate (ZHA) | 0.169 | 0.333 | -0.163 | 0.872 | **-0.044** | -0.058 | -0.031 | 0.380 | -0.112 | 0.924 |
| 1-Butene-4-isothiocyante (ITC) | 0.595 | 0.334 | -0.607 | 1.313 | **-0.202** | -0.323 | -0.132 | 0.575 | -0.434 | 1.529 |

| | Observed changes | Predicted changes | | | | | | | | |
|---|---|---|---|---|---|---|---|---|---|---|
| | | Total | 95% HPD | | Direct | 95% HPD | | Indirect | 95% HPD | |
| | | | lower | upper | | lower | upper | | lower | upper |
| ***(B) Hoverfly*** | | | | | | | | | | |
| Plant height (Height) | -0.292 | **0.499** | 0.320 | 0.660 | **0.549** | 0.368 | 0.695 | -0.054 | -0.162 | 0.016 |
| Petal width (PW) | 0.154 | 0.047 | -0.055 | 0.198 | **-0.055** | -0.073 | -0.042 | 0.124 | -0.004 | 0.255 |
| Petal length (PL) | 0.391 | **0.160** | 0.046 | 0.308 | **0.029** | 0.022 | 0.038 | **0.135** | 0.024 | 0.282 |
| Flower diameter (FD) | **0.254** | **0.298** | 0.143 | 0.460 | **0.139** | 0.101 | 0.203 | **0.134** | 0.008 | 0.284 |
| Benzaldehyde (Ben) | 0.645 | -0.055 | -0.189 | 0.083 | **-0.081** | -0.109 | -0.061 | 0.024 | -0.109 | 0.163 |
| Phenylacetaldehyde (PAA) | 0.373 | -0.012 | -0.193 | 0.128 | **-0.145** | -0.225 | -0.096 | 0.114 | -0.029 | 0.292 |
| α-Farnesene (FAR) | 0.746 | 0.021 | -0.123 | 0.132 | **-0.102** | -0.142 | -0.082 | 0.105 | -0.007 | 0.258 |
| Benzyl nitrile (BenN) | 0.637 | 0.018 | -0.096 | 0.193 | **0.102** | 0.072 | 0.134 | -0.072 | -0.202 | 0.100 |
| 2-Amino benzaldehyde (Aben) | -0.009 | 0.040 | -0.119 | 0.192 | **-0.025** | -0.033 | -0.014 | 0.088 | -0.094 | 0.217 |
| Indole (Ind) | **0.041** | **0.140** | 0.017 | 0.331 | **0.125** | 0.086 | 0.175 | 0.025 | -0.121 | 0.187 |
| Methyl anthranilate (MeA) | 0.257 | -0.043 | -0.240 | 0.091 | **-0.213** | -0.304 | -0.135 | 0.145 | -0.031 | 0.291 |
| Phenylethyl alcohol (PA) | 0.396 | 0.023 | -0.132 | 0.187 | **-0.012** | -0.019 | -0.009 | 0.035 | -0.123 | 0.196 |
| Methyl salicylate (MeS) | -1.078 | 0.012 | -0.170 | 0.128 | **0.064** | 0.044 | 0.089 | -0.062 | -0.243 | 0.072 |
| Methyl benzoate (MeB) | 0.744 | -0.027 | -0.187 | 0.097 | **-0.056** | -0.076 | -0.040 | 0.041 | -0.127 | 0.156 |
| Z-(3)-Hexenyl acetate (ZHA) | -0.707 | -0.071 | -0.201 | 0.070 | **-0.112** | -0.148 | -0.080 | 0.037 | -0.080 | 0.187 |
| 1-Butene-4-isothiocyante (ITC) | 0.798 | 0.064 | -0.171 | 0.193 | **-0.003** | -0.006 | -0.002 | 0.006 | -0.167 | 0.197 |

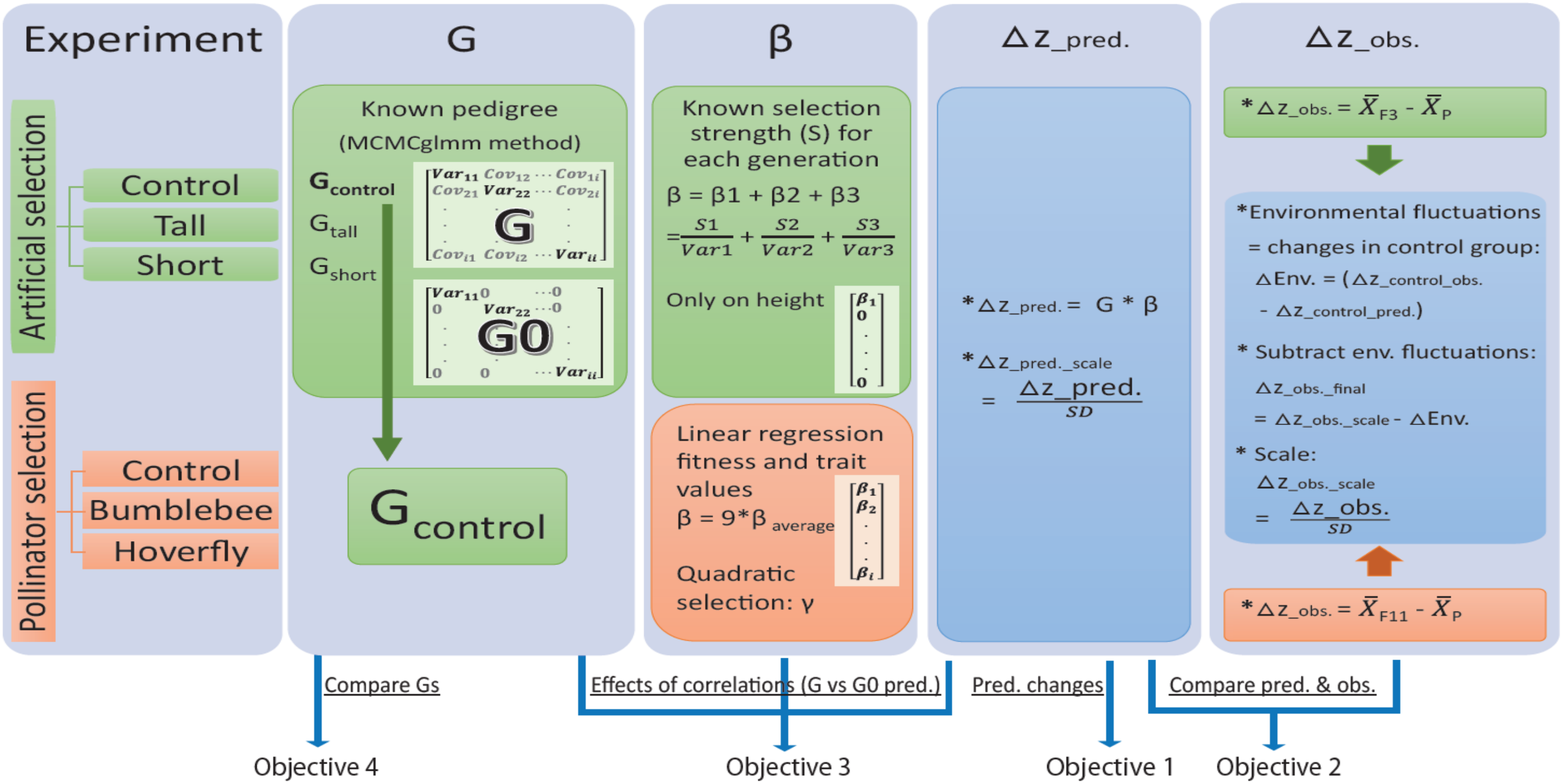


**Figure S1.** Workflow of the main analysis procedures and the study objectives.
